# Direct delivery of Cas9 or base editor protein and guide RNA complex enables genome editing in the retina

**DOI:** 10.1101/2023.10.16.562239

**Authors:** Catherine Botto, Juliette Pulman, Hugo Malki, Duohao Ren, Paul Oudin, Anne De Cian, Marie As, Charlotte Izabelle, Bruno Saubamea, Stéphane Fouquet, Camille Robert, Aziz El-Amraoui, Sylvain Fisson, Jean-Paul Concordet, Deniz Dalkara

**Affiliations:** Sorbonne Université, INSERM, CNRS, Institut de la Vision, 17 rue Moreau, F-75012 Paris, France; Université Paris-Saclay, Univ Evry, Inserm, Genethon, Integrare Research Unit UMR_S951, 91000 Evry-Courcouronnes, France; Laboratoire Structure et Instabilité des Génomes - INSERM U1154 - CNRS 7196, Muséum National d’Histoire Naturelle - CP26 43 rue Cuvier 75231 Paris Cedex 05; Université Paris Cité, Inserm, CNRS, P-MIM, PICMO, F-75006 Paris, France; Institut Pasteur, Université Paris Cité, INSERM AO06, Institut de l’Audition, Unit Progressive Sensory Disorders, Pathophysiology and Therapy, 63 rue de Charenton F-75012 Paris, France

## Abstract

Genome editing by CRISPR-Cas holds promise for the treatment of retinal dystrophies. For therapeutic gene editing, transient delivery of CRISPR- Cas9 is preferable to viral delivery which leads to long-term expression with potential adverse consequences. Successful delivery of Cas9 protein and its guide RNA as ribonucleoprotein (RNP) complexes has been reported in the retinal pigment epithelium *in vivo* but not into photoreceptors, the main target of retinal dystrophies. Here, we investigate the feasibility of direct RNP delivery to photoreceptors and RPE cells. We show that RNPs composed of Cas9 or adenine-base editor and guide RNA, without addition of any carrier compounds, induce gene editing in retinal cells at variable rates depending on the guide RNA efficiency and on the locus. But Cas9 RNP delivery at high concentrations leads to outer retinal toxicity indicating a need to improve delivery efficiency for future therapeutic use.

## Introduction

Gene supplementation has been successful for the treatment of rare inherited retinal degenerations (IRD) caused by single recessive mutations on genes small enough to be delivered via adeno-associated virus (AAV). Allele-specific genomic ablation or gene correction using gene editing may extend these outcomes to diseases caused by dominant mutations or by mutations on large genes, which altogether affect a larger group of patients (*1*). To date, the most advanced gene editing application in the eye is designed to excise a cryptic splice site in the *CEP290* gene to improve vision in patients suffering from Leber congenital amaurosis type 10 (LCA10). This strategy based on subretinal delivery of AAV encoding SaCas9 and two guide RNAs (sgRNAs) has been successful in non-human primates (*2*) and has served as the basis for the first clinical trial of gene editing in the eye (NCT03872479).

Unfortunately, viral mediated delivery of CRISPR-Cas9 gene editing reagents raises the possibility of permanent integration in the genome (*3*), off-target gene disruption through long-term exposure to gene editing reagents as well as potential immune reactions to these proteins of bacterial origin (*4*). In addition, 4.7 kb carrying capacity of AAV limits the use of new gene editing tools such as base and prime editors that can be used to treat ups to 99.9% of IRD (*5–7*). In this context, transient delivery of Cas9 protein and its sgRNA as ribonucleoprotein (Cas9 RNP) using non-viral vectors is an attractive alternative for CRISPR-Cas9- mediated gene editing as its rapid degradation might limit its off-target effects and its non-DNA form will rule out the risk of genome integration. The first attempt to Cas9 RNP delivery in the retina employed cationic lipids and reached 22% editing efficiency in the gene encoding Vascular Endothelial Growth Factor A (*Vegfa)* in the retinal pigment epithelium (RPE). These results were obtained in an acute mouse model of age-related macular degeneration (*8*). A second study confirmed the capacity of lipid vectors to deliver Cas9 RNP to the RPE, but in wild-type mice, resulting in a more modest indel rate of 6% at the same locus in isolated EGFP positive RPE cells. This study also reported signs of toxicity at high Cas9 RNP concentrations (*9*). Several other studies using lipid nanoparticles or nanocapsules have followed work on RNP delivery to the RPE with similar indel rates (*10*, *11*). Although these first results of gene editing in the RPE using Cas9 RNP are promising, so far, no study has reported Cas9 RNP mediated gene editing in cells of the neural retina, notably in the photoreceptors. As the majority of mutated genes in IRD are expressed in photoreceptor cells, these cells are likely to be the main targets of gene editing applications in the coming years. To fill this unmet need, we investigated transient delivery of Cas9 protein and base editor as RNP complexes in the neural retina in comparison to RPE cells. We tested different categories of non-viral vectors that display different physico-chemical properties to assess their ability to complex and deliver Cas9 RNP into retinal cells. We show that without any vector, Cas9 RNPs induce up to 15 % of indels in the whole neural retina of wildtype adult mice. Similarly, base editor RNPs without addition of any carrier compounds trigger up to 17 % gene editing in the neural retina when injected without any carrier compounds. The editing efficiency is dependent on the dose of RNP, the identity of the sgRNA, and the level of expression of the targeted genes in the given cell population. We also show that physical barriers specific to our tissue of interest include photoreceptor outer segments and outer limiting membrane, which impede the entry of the RNPs into the neural retina. The delivery efficacy is further hampered by intracellular barriers, such as nuclear entry and the accessibility of the targeted gene within the genome once the RNP gets into the nucleus. Altogether our work highlights the key parameters to consider in improving Cas9 RNPs or designing new classes of carrier compounds to bring RNPs into the retina for therapeutic gene editing. Moreover, it highlights the cell and gene specific features of Cas9 mediated gene editing that must be considered in the design and preclinical testing of gene editing therapeutics.

## Results

### Ribonucleoprotein complexes (Cas9 protein and its sgRNA) induce indels in the neural retina and in the RPE

Prior to in vivo experiments, we first examined the properties of Cas9 protein alone or coupled with its sgRNA (ribonucleoprotein -RNP- complex) using transmission electron microscopy (TEM) and dynamic light scattering (DLS). We used a Cas9 from *Streptococcus Pyogenes* (SpCas9; net charge +22) with two nuclear localization signals (NLS, net charge +5). The protein alone was visible using TEM, displaying a size smaller than 10 nm (Fig. 1A). We then complexed it with a previously described sgRNA targeting *Vegfa* gene (net charge -120) (*8*, *9*). The complexation of the SpCas9 proteins with two NLS tags and this sgRNA gave rise to homogenous RNP complexes (theorical net charge of -88) with an average size of 17 nm (Fig 1A). These observations were confirmed by dynamic light scattering (DLS) showing a particle size distribution between 10 and 70 nm with a peak of ∼24 nm, consistent with previous reports (*12*) (Fig. 1B).

**Figure 1.**
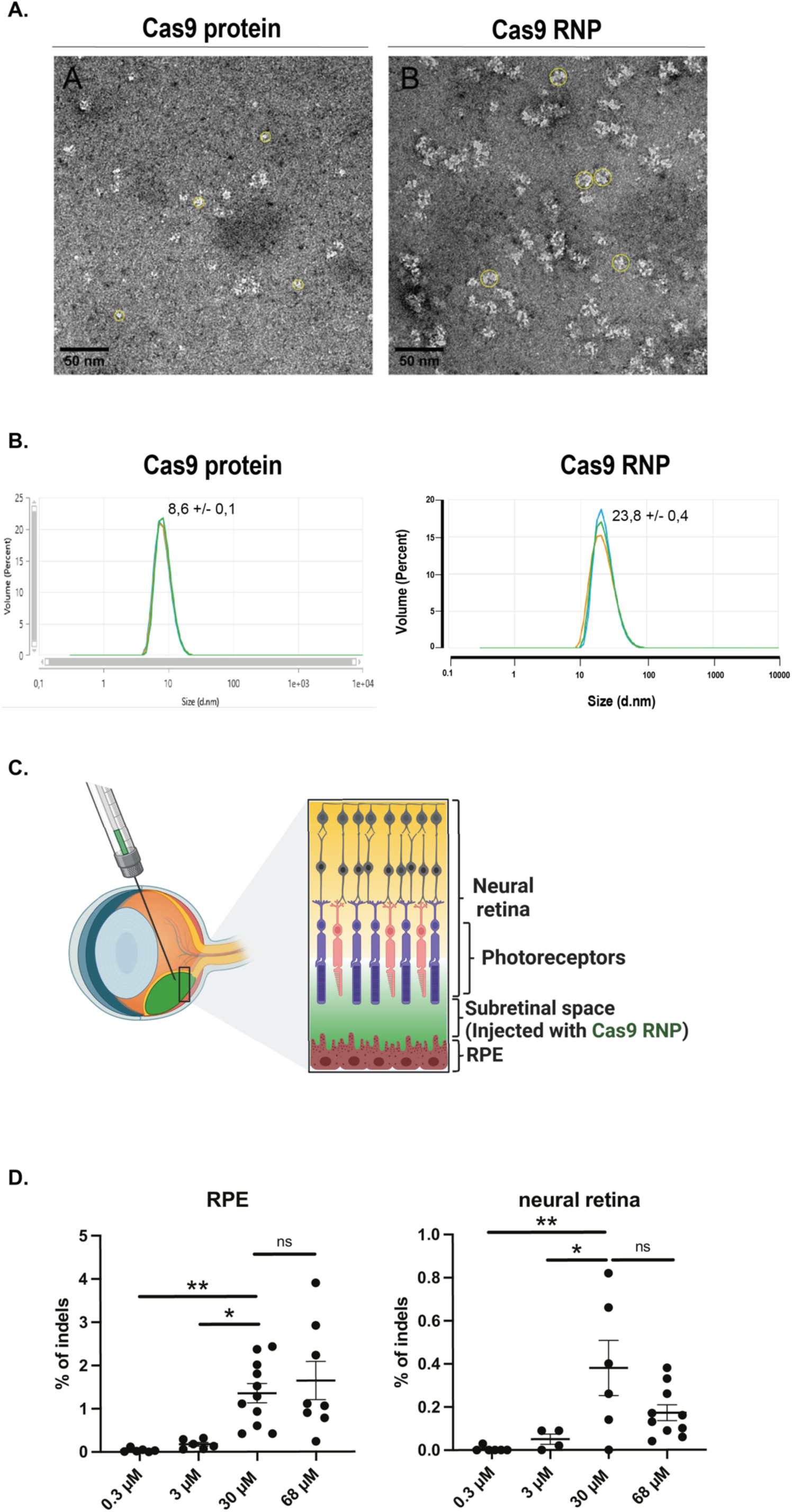
Direct Cas9 RNP delivery induces indels *in vivo* in the neural retina and in the RPE. **(A)** Transmission electron microscopy (TEM) analysis of 60 µM Cas9 protein (10 nm diameter circles are shown in yellow) and Cas9 RNP (17 nm diameter circles are shown in yellow) imaged using 1% aqueous uranyl acetate as a negative stain. **(B)** Size (nm) of 30 µM Cas9 protein alone or Cas9 RNPs determined by dynamic light scattering (DLS). Three different measures (in blue, red and green) were performed at 5-minute intervals. **(C)** Schematic representation of the sub-retinal injections performed in the eyes of adult wild type mice. RPE and neural retinas are separated during dissections and analyzed independently. **(D)** Indels in the RPE and neural retina after sub-retinal injection of Cas9 RNPs at different concentrations *in vivo*. NGS analysis was performed 7 days after injection. Mean ± SEM. Ordinary One-way ANOVA test, Dunnett’s multiple comparisons test.

Next, we injected these RNPs subretinally without any vector in adult wild-type mice. One week after injection, DNA was extracted from the whole RPE and the whole neural retina of each eye and indels were quantified at the targeted *Vegfa* locus using next generation sequencing (NGS) (Fig. 1C). A dose dependent increase in indels was observed both in the RPE and neural retina up to 30 µM RNP, with no improvement at higher doses (Fig. 1D). The dose of 30 µM was therefore chosen all follow-up experiments. Interestingly, a higher indel rate of 1.4 % was obtained in the RPE, as compared to 0,4% indels obtained in the neural retina reaching at the 30 µM dose (Fig. 1D).

### Limited diffusion of Cas9 proteins into the neural retina

To understand the low activity of the Cas9 RNPs in the neural retina, we examined their distribution three days post-injection. 3D imaging of the whole eye allows the detection of the Cas9 protein. After segmentation of the neural retina, we see that Cas9 proteins are still present in clusters in some areas (Fig. 2A and Movie S1). We also see that, in Cas9 RNP injected eye, within the bleb, some areas within the bleb are completely devoid of Cas9, suggesting that Cas9 has already been partially degraded (Fig. 2A and Movie S1).

**Figure 2.**
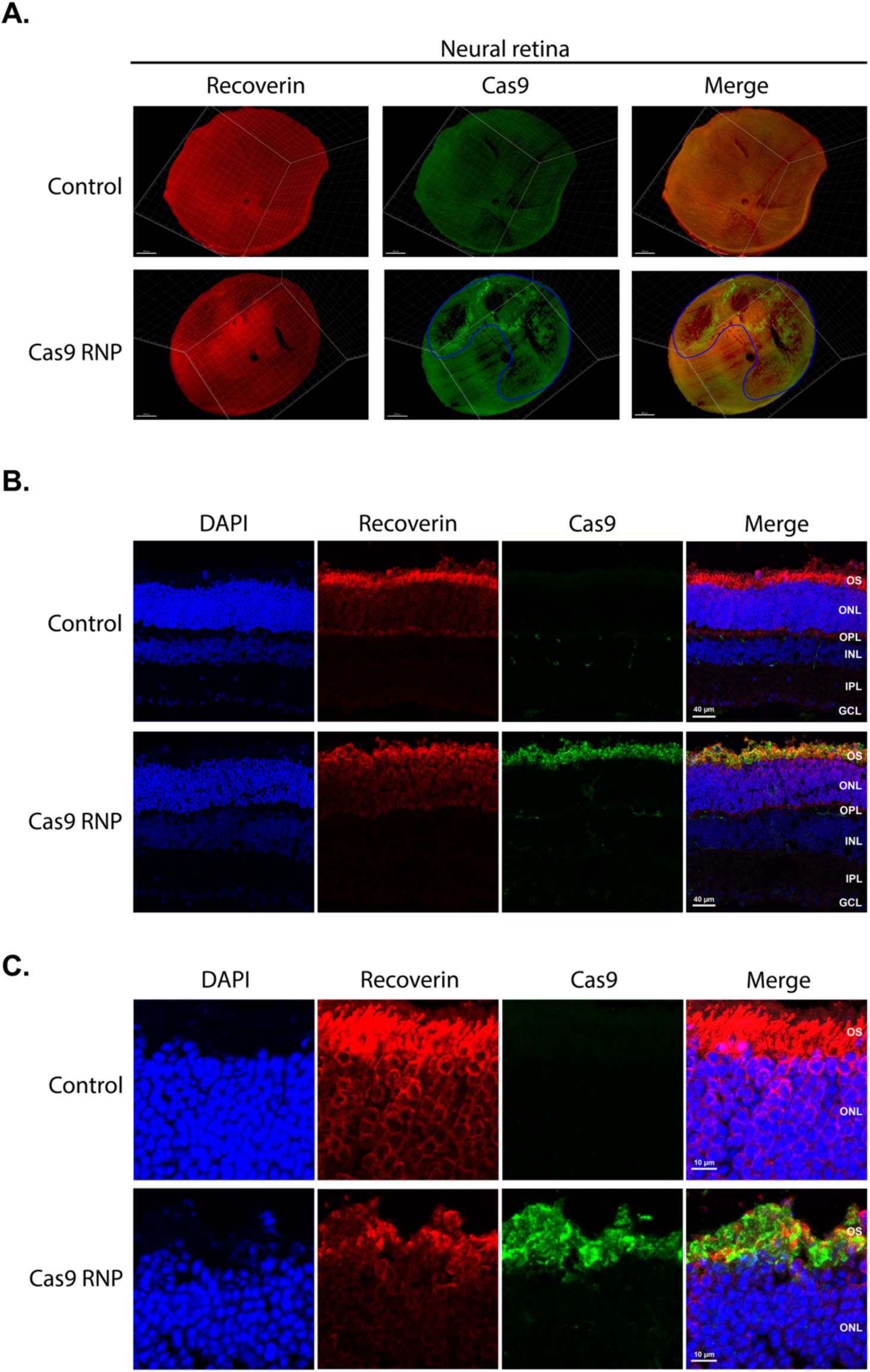
Localization of Cas9 protein inside the neural retina. Eyes were collected three days after 2 µL subretinal injection of buffer (control) or 30 µM Cas9 RNPs. (A) Top view of the entire neural retina after segmentation from mouse whole eye after clearing and 3D imaging. Injection zone area is circled in blue. Staining of the photoreceptors (Recoverin, red) and Cas9 protein (green) (B-C) Neural retina cross-sections were stained for nuclei (DAPI, blue), photoreceptors (Recoverin, red) and Cas9 protein (green). OS: outer segment, ONL: outer nuclear layer, OPL: outer plexiform layer, INL: inner nuclear layer, IPL: inner plexiform layer, GCL: ganglion cell layer (C) Zoom on the photoreceptors outer segment and nuclei.

To have a better resolution of the different layer of the neural retina, we performed immunohistochemistry against the Cas9 protein on retinal cryosections. We see that the majority of Cas9 proteins was not able to infiltrate into the neural retina (Fig. 2B). When zooming on the photoreceptor cells, we see that specific barriers, such as outer segments discs, external limiting membrane and tight extracellular matrix of the outer limiting membrane, limit the diffusion of RNPs and that Cas9 RNP disorganized the OS (Fig. 2C).

### Cationic lipids didn’t improve Cas9 RNP delivery to the retina

We next investigated if physical barriers specific to the neural retina could be bypassed by using cationic lipids as non-viral vectors. Cas9 RNP entry into cells may not be favored by the negative global charge of these complexes (*13*). To enter cells by endocytosis or direct penetration through the cell membrane after interaction with proteoglycans, positive surface charges are preferred. Based on previous studies in the inner ear and in the RPE (*8*, *13*), we investigated the capacity of the commercial cationic lipids Lipofectamine 2000 or Lipofectamine RNAiMAX to deliver Cas9 RNPs to neural retinal cells. However, Cas9 RNPs mixed with Lipofectamine 2000 gave heterogenous assemblies of complexes, forming either uncomplexed monomeric RNPs and liposomes or aggregates and multilamellar lipoplexes of different sizes (Fig. 3A-H). DLS measurements confirmed heterogenous composition of the mix with peaks corresponding to the detection of aggregates (Fig. 3I and Fig. S1). Finally, after sub-retinal injection in wild-type mice, neither Lipofectamine 2000 nor RNAiMax improved RNP delivery in the retina (Fig. 3J).

**Figure 3.**
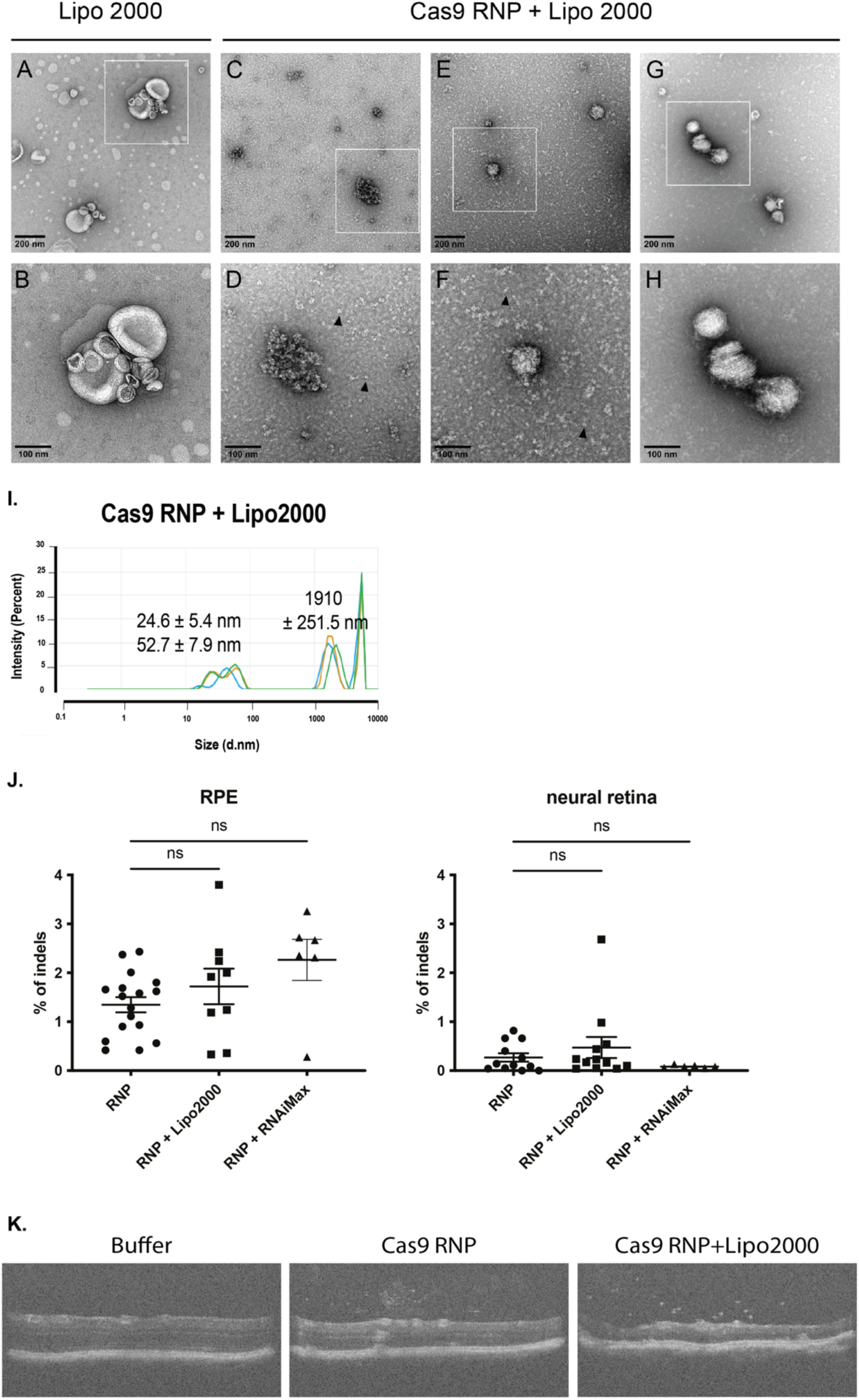
RNP delivery to the retina using cationic lipids: heterogenous assemblies of complexes, no efficacy improvement, and toxicity. **(A to H)** TEM analysis of (A-B) Lipofectamine 2000 undiluted and (C-H) different complexes observed when mixing Cas9 RNP and Lipofectamine 2000 (diluted 100X). Monomers of RNP (D and F) are highlighted with black arrows. **(I)** Size (nm) of 30 µM Cas9 RNP complexed with Lipofectamine 2000 determined by DLS. Three different measures (in blue, red and green) were performed at 5-minute intervals. **(J)** Indels induced in the whole RPE and neural retina after 2 µL of sub-retinal injection of 30 µM Cas9 RNPs complexed with cationic lipids: Lipofectamine 2000 or RNAiMAX *in vivo* in WT mice. NGS analysis was performed 7 days post-injection. Mean ± SEM. Ordinary One-way ANOVA test, Dunnett’s multiple comparisons test. **(K)** OCT images 1-month post-injection of buffer solution, 30 µM Cas9 RNPs or Cas9 RNPs complexed with lipofectamine 2000 in adult WT mice.

As Cas9 RNP delivery with lipoplexes showed sign of toxicity in the RPE (*9*), we also investigated the potential toxicity of our naked RNP on the neural retina of WT mice by *in vivo* imaging. Optical coherence tomography (OCT) was performed 1 month after Cas9 RNP sub-retinal injection and showed retinal thinning at the injection site and cellular infiltrates in eyes injected with RNP alone or RNP mixed with Lipofectamine 2000 (Fig. 3K). In eyes injected with Cas9 RNP alone, retinas were less damaged than those injected with Cas9 RNP mixed with Lipofectamine 2000 (Fig. 3K).

We next switched to a peptide-based system aiming to improve the delivery of RNPs to the neural retina. A previous study had revealed the capacity of a shuttle peptide, named SP10, to promote complexation and delivery of Cas9 RNP into epithelial cells *in vivo* (*14*). SP10 is composed of a cell-penetrating peptide (CPP) linked to an endosomolytic peptide for endosome rupture (net positive charge + 10). However, mixing of 250 µM SP10 and 30 µM Cas9 RNPs resulted in heterogenous particles of 34.4 ± 3.7 nm and aggregates (Fig. S3) and failed to significantly increase the capacity of RNPs to induce indels in the retina (Fig. S3).

### Editing efficiency partially depends on the expression level of the targeted genes but mostly depends on the gRNA efficiency

After investigating extracellular barriers to RNP delivery to the retina, we sought to identify intracellular factors contributing to gene editing efficiency in retinal cells. To better understand the contribution of chromatin accessibility on editing efficiency in photoreceptors, we compared indel efficiencies obtained with our *Vegfa* sgRNA, highly expressed in the RPE, to other sgRNAs targeting genes highly expressed in the photoreceptors (Fig. S4). It is important to note that genes highly expressed in photoreceptors are often mutated in monogenic retinal diseases, and therefore constitute major targets for future therapeutic gene editing applications (*1*). *Sag*, coding for S-arrestin protein; *Rho,* coding for Rhodopsin; and *Pde6b* genes were selected and sgRNAs were designed in silico. The efficacy of each guide was assessed in a murine cone cell line using the TIDE assay and compared to the previously tested guide targeting *Vegf*a gene. Among sgRNAs for *Sag* gene, sgRNA 3 in exon 8 induced the highest rate of indels, with 24,5% of indels, which was comparable to that induced by the *Vegfa* sgRNA (Fig. 4A). Focusing on the best sgRNA targeting each of the *Sag, Rho and Pde6b* genes, we tested further their efficacy *in vivo*. In genes expressed in the photoreceptors, we observed an increase in the indel rates in the neural retina compared to the RPE (Fig. 4B). On the contrary, when targeting *Vegfa*, which is not expressed in the photoreceptors, the rate of indels was lower in the neural retina than in the RPE (Fig. 4B). These results suggest that the editing efficiency depends on the level of expression of the target genes. As Sag gRNA3 show a massive increase in the indels (7,2% ± 3,9%) in the neural retina compared with all the others, we wondered if this was specific of the gene availability or of the gRNA. So, we tested the two other sgRNAs screened *in vitro* for Sag. Interestingly, sgRNA 1 and 2 gave very low indels (Fig. 4C), suggesting that the efficacy highly depends on the gRNA sequence and the position on the targeted gene locus. This suggests that therapeutic gene editing might be more successful in certain gene targets than others. Furthermore, this significant difference in indel efficacy of *Sag* sgRNA3 was not visible during the *in vitro* screening (Fig. 4A) suggesting gRNA screens in vitro have only partial predictive value of *in vivo* efficacy. Finally, these results shows that Cas9 RNPs can efficiently edit photoreceptors when combined with a sgRNA targeting a transcriptionally active locus.

**Figure 4.**
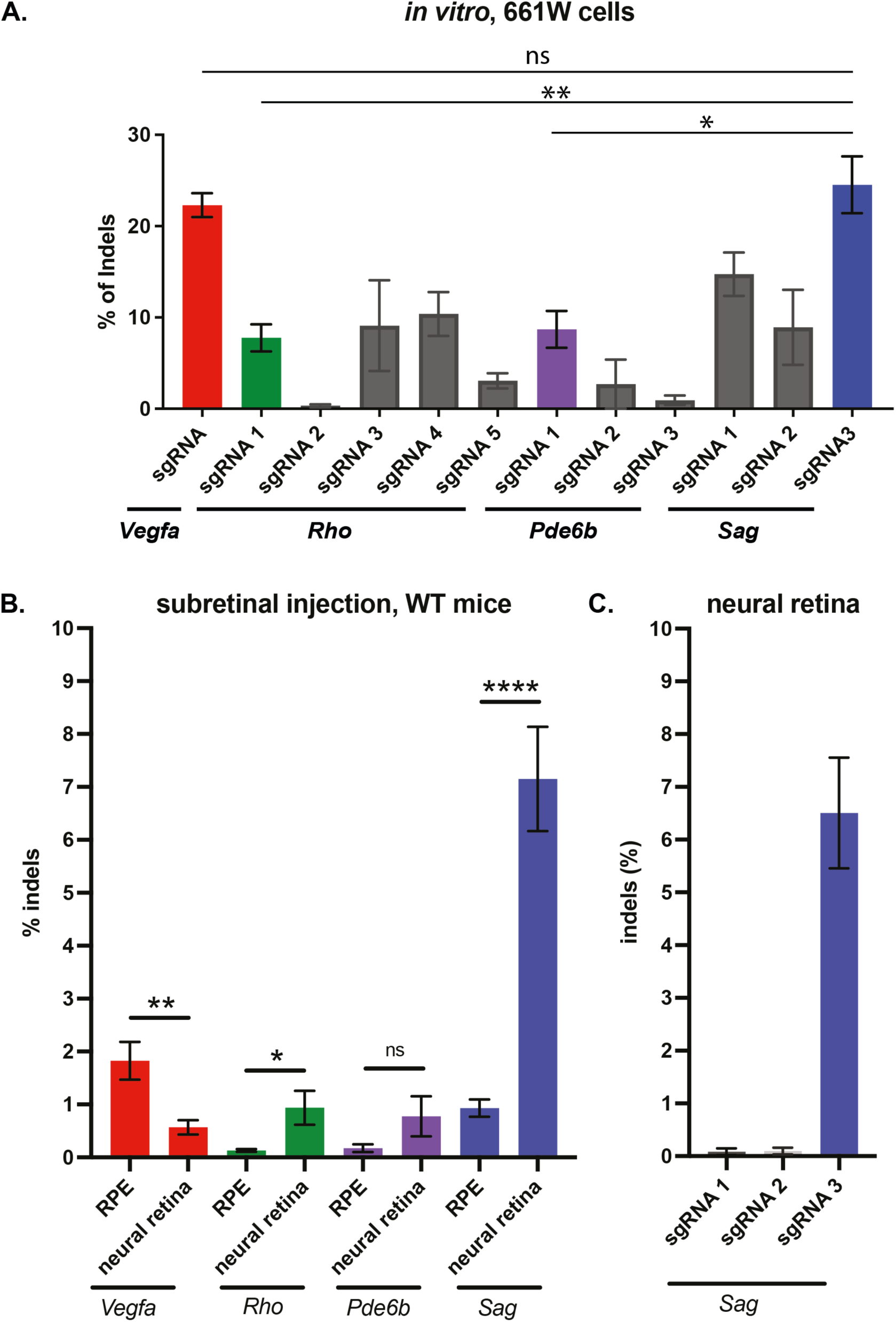
Efficiency of different sgRNAs targeting genes highly expressed in the retina. **(A)** Validation of sgRNAs targeting genes highly expressed in photoreceptors (*Sag*, *Rho* and *Pde6b*), compared with the sgRNA targeting *Vegfa* gene by TIDE assay. Transfection of Cas9 RNPs complexed with lipofectamine 2000 in 661W cell line. Mean ± SEM. Ordinary One-way ANOVA test, Dunnett’s multiple comparisons test. sgRNAs that were selected for further *in vivo* studies are highlighted in colors. **(B)** Frequencies of indels induced in the whole RPE and whole neural retina of WT mice after sub-retinal injection of 30 µM Cas9 RNPs with *Sag* sgRNA3; *Rho* sgRNA1 or *Pde6b* sgRNA1 and compared with the *Vegfa* sgRNA. NGS analysis was performed 7 days post-injection. Mean ± SEM. Ordinary One-way ANOVA test, Dunnett’s multiple comparisons test. **(C)** Frequencies of indels induced in the whole neural retina of WT mice after sub-retinal injection of 30 µM Cas9 RNPs with *Sag* sgRNA1; *Sag* sgRNA2 or *Sag* sgRNA3. NGS analysis was performed 7 days post-injection. Mean ± SEM.

### Base editors RNP can also induce indels in the neural retina and in the RPE

One significative advantage of direct RNP delivery over AAV mediated delivery of Cas9 and gRNA is that there is theoretically no size limit for this mode of delivery: nucleases, base editors, transposases/recombinases and prime editors could all be delivered without any carrier vector, only complexed with their sgRNAs. To prove this point and to test under identical conditions the efficacy of base editing compared to Cas9, we evaluated the editing efficacy of naked base editor RNPs complexed to the Sag sgRNA3. Our adenine base editor proteins alone were detected at 86.6 nm, suggesting a possible aggregation of particles when uncomplexed, while our complexation of with gRNA gave particles of 43.4 nm with few aggregates at 2,6 µm (Fig 5A). When injected subretinally, the base editor RNPs showed a similar editing efficacy than the Cas9 RNP, yielding 10,3% ± 5,1% of targeted base changes (Fig 5B), with 0,2% of bystander effect and no detectable indels. We also confirmed the presence of the targeted substitution in the cDNA with a significant correlation with the substitution found at the DNA level (Fig 5C). These results show that RNP delivery is feasible across different proteins and that robust outcomes can be obtained in terms of gene editing efficacy when targeting the same locus. This result further shows that RNP delivery can be an advantageous mechanism to do side-by-side comparisons of genome editing reagents.

**Figure 5.**
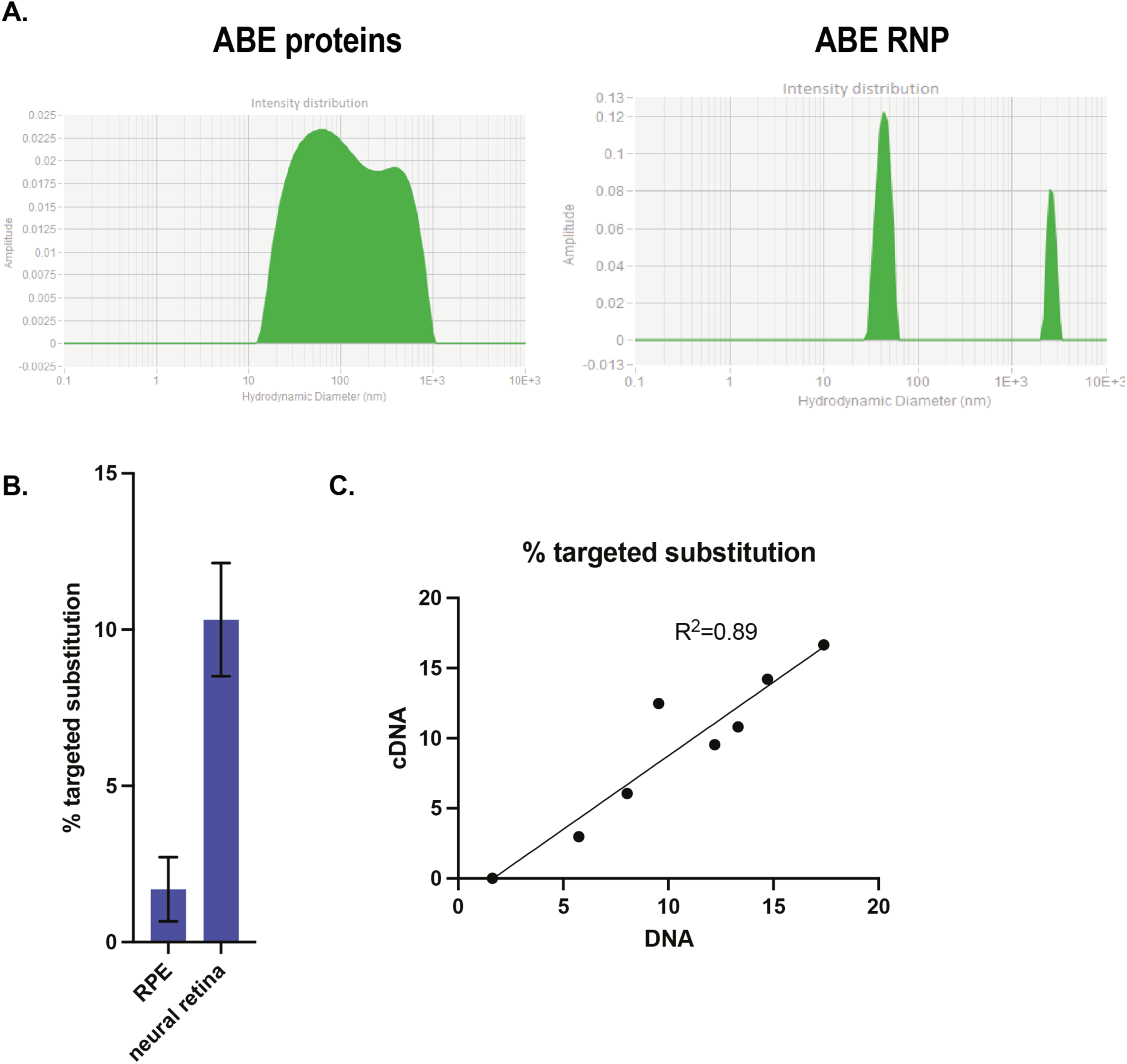
ABE RNP delivery to the retina is able to generate targeted substitution. **(A)** Size (nm) of 30 µM ABE proteins alone or ABE RNP determined by DLS. **(B)** Frequencies of on-site substitutions in the RPE and the whole neural retina of WT mice after sub-retinal injection of 30 µM ABE RNPs with *Sag* sgRNA3. NGS analysis was performed 7 days post-injection. Mean ± SEM. Student’s t-test. **(C)** Correlation between the targeted substitution found at the DNA level and at the cDNA level by NGS sequencing performed 7 days post injection.

## Discussion

Regardless of the gene’s size and nature of the disease-causing mutation, gene editing tools provide unprecedented opportunities for effective and long-lasting treatment of both dominant and recessive forms of vision loss (*5*, *15*). In this context, persistent expression of the gene editing proteins is undesirable and there is high interest in developing a transient delivery system for their safe use in gene therapy. To this aim, we investigated how to deliver Cas9 and its sgRNA as a ribonucleoprotein into retinal cells with a particular focus on the neural retina *in vivo*. We showed that Cas9 RNPs, injected into the subretinal space without any carrier compounds generate up to 15% of indels on the *Sag* gene in the neural retina of adult wild-type mice. To our knowledge, this is the first report of an editing efficiency for an RNP molecule delivered without any vectors in the RPE and the neural retina. This may have therapeutic relevance in some diseases. Indeed, Maeder and colleagues estimated an editing efficacy of 10% may provide measurable benefit to LCA10 patients affected by mutations in *CEP290* (*2*). However, the percentage of gene edition needed for a therapeutic effect will highly depend on the disease-causing gene, pathogenic mechanisms and kinetics.

Until now, Cas9 RNPs had shown efficiency only in the RPE and not in the neural retina. The gene editing efficacy in the RPE varied between 6 to 24% within the injected area following subretinal delivery(*8*, *9*). The variability might have been the result of different animal models being used alongside methodological differences between studies. In our case, instead of analyzing indel rates in the injected area (covering around half of the retinal surface only); we measured the indel rates of the entire RPE and neural retina. Under these conditions, we achieved 7-10% of indels in the whole neural retina and 2% in the RPE and show for the first time that RNPs delivered alone can edit photoreceptor cells in the neural retina. In comparison, in the neural retina, AAV delivered Cas9 and sgRNAs targeting various loci (*1*). For example, Wu et al. used a dual AAV8 vector with saCas9 and a double sgRNA achieving 17,8% of indels in the whole neural retina (*16*). Editing efficiency of our RNP appears lower compared to this study using AAV but this observed difference could be due to the use of different models, differences in sgRNA efficacy, and the use of two guide RNAs.

As regards to base editing, BEs have been delivered by subretinal injection via a dual AAV into photoreceptor cells, resulting in 21 to 26% editing of the photoreceptor cells (*17*, *18*). BEs were also delivered using a lentiviral vectors, into the RPE resulting in around 16% of editing rate (*19*). Finally, BEs have been reported to be delivered as RNP complexed with Lipofectamine 2000 via subretinal injection in adult mice eyes and yielded 2% efficiency in RPE cells (*20*). Our findings show that naked RNPs achieved five times more base editing in the whole neural retina. Although at this stage, gene editing in the retina using RNP still is half as efficient as using AAVs.

The use of a non-viral vectors is likely necessary to improve the delivery efficacy, and lower the dose (and hence toxicity) of RNPs needed for therapeutic gene editing. Cationic lipids such as Lipofectamine 2000 or peptide-based carriers used in this work didn’t improve the editing efficiency and were highly toxic for the retina suggesting other types of vectors or conjugates that are tailored to retinal cells are needed. But RNPs alone also induced toxicity. These findings further confirm that a decrease in Cas9 concentration and its potential shielding from immune cells in subretinal space are needed. We demonstrated that following RNP injections into the subretinal space, the majority of Cas9 proteins accumulate over the neural retina and seem unable to infiltrate the dense extracellular matrix of the outer nuclear layer. Photoreceptors form a dense layer of cells that present specific barriers such as photoreceptor outer segments and the external limiting membrane. Targeting elements, increasing the capacity of the complexes to bind and enter photoreceptor cells, will be crucial to increasing delivery efficacy (*11*, *21*). Attempts to decorate RNP nanocarriers with molecules like all-trans retinoic acid have been successful to increase entry into the RPE but so far there have not been any reports of compounds increasing targeting to photoreceptors (*11*).

Once infiltration across the extracellular physical barriers and intracellular entry is obtained, it is also of great interest to increase the endosomal escape capacity of RNPs as only the complexes that escape ubiquitination and degradation have a chance to reach the nucleus (*22*). Nuclear entry can be improved to increase chances of successful gene editing using RNPs. Addition of NLS sequences has previously been explored towards this aim showing that only a small number of NLS are beneficial as too many NLS sequences sterically hinder binding to the target DNA locus (*23*). In our study, we use a Cas9 RNP with 2 NLS.

Then, as RNP complexes reach the nucleus, the role of the sgRNA seems to be the most important parameter determining gene editing rates. We demonstrate that the sgRNA efficacy varies depending on the cell type and the level of expression of the targeted gene. Indeed, our editing rates were higher when targeting photoreceptor expressed genes in photoreceptors and vice versa in the RPE. This is likely due to the accessibility of the chromatin to the RNP complex and the DNA reparation system (*24*). But we showed that efficiency mostly depends on the sgRNA sequence and the position on the targeted gene locus. The sgRNA for a specific experiment is currently chosen by applying design tools that predict the most active and efficient guides using machine learning but these tools do not take into account cell-type specific features such as chromatin accessibility and cell cycle state of the cell. The computational scoring methods place emphasis on minimizing off-target binding by minimizing mismatches with the target sequence. However other parameters of the sgRNA, such as the GC content, of the sgRNA will affect its stability and therefore the editing efficiency. Selecting the right design tool, and experimental workflow, achieving a successful gene-editing experiment is now within the capabilities of almost every biological laboratory for *in vitro* applications. But *in vivo* gene editing is tissue and cell-type specific, requiring additional experiments to extend *in vitro* considerations.

We also show that RNP delivery is feasible across different proteins, with similar results between Cas9 RNP and base adenine base editors. The RNP delivery strategy described here can be of use for quick side-by-side comparison in such a context.

In conclusion, our work shows the possibility to transiently deliver Cas9 or base editors directly as an RNP into the RPE and the neural retina *in vivo* and identifies the set of parameters for optimization of RNP delivery strategies into this tissue.

## Material and Methods

### sgRNA design

sgRNAs targeting mouse *Pde6b*, *Sag* and *Rho* genes were designed using CRISPOR outside of nucleosome regions predicted with (*25*, *26*). The sgRNA the vascular endothelial growth factor A (*Vegfa*) gene was published by Kim and colleagues (*8*). All sgRNAs were synthesized and purified using the GeneArt^TM^ Precision gRNA Synthesis Kit (Invitrogen), according to the manufacturer’s protocol. sgRNAs eluted in water were aliquoted and stored at -80°C. All 20pb sequences targeted by each sgRNA are listed in supplementary Table S2.

### Cas9 nuclease

For in vivo experiments, Streptococcus pyogenes Cas9 (SpCas9) nuclease with 2 nuclear localization sequences (NLS) (one on its N and one on its C terminal) was produced as previously described in Menoret et al (*27*) and kept at -80°C until use. For in vitro experiments, SpCas9 nuclease (Aldevron) was aliquoted and kept at -20°C until use. Plasmid for E.Coli expression of ABE (Addgene #161788 from the Liu lab) was modified to express the ABE variant ABE8-13m (*28*) and ABE8- 13m purification was performed as recommended in Huang et al (*29*).

### RNP preparation and complexation

Ribonucleoproteins (RNPs) were prepared immediately before use. Briefly, SpCas9 proteins were mixed to sgRNAs at a molar ratio Cas9 protein/sgRNA of 1:1 in a final buffer concentration of 20 mM HEPES/200 mM KCL (pH7.4). SpCas9/sgRNA solution was incubated at room temperature for 5 minutes before direct use or complexation with a vector. Freshly prepared RNPs were mixed with several non-viral vectors, then solutions were vortexed for 10 seconds and incubated at room temperature for 10 minutes before direct use. Lipofectamine™ 2000 and Lipofectamine**™** RNAiMAX Transfection Reagent (Invitrogen) were purchased from ThermoFischer. Shuttle peptides were synthetized by Covalab.

### Transmission electron microscopy (TEM)

Samples were prepared following the negative staining protocol reported by (*30*). Briefly, a glow discharged Carbon/Formvar grid (Agar Scientific, Stansted, United Kingdom) was inverted onto a 5 µL drop of the sample. After 1 minute, the grid was blotted with filter paper and rinsed by quickly touching a drop of water and blotting (three times). The grid was then floated consecutively on three drops of 1% aqueous uranyl acetate for 10 sec, 10 sec and 1 min, blotted and air dried for 20 minutes before observation. Images were acquired using a Jeol 1400 Flash Transmission Electron Microscope (Jeol, Croissy-sur-Seine, France) operated at 120 kV and equipped with a RIO CMOS camera (Ametek SAS, Elancourt, France).

### Dynamic Light Scattering (DLS)

The hydrodynamic diameter (size) of each complex was measured using DLS (Zetasizer nano analyser ZS; Malvern Instruments) at Paris ESPCI facility or using the Stunner (Unchained Labs). Three different measures were carried out with 5 minutes in between each.

### Cell culture and transfection

Murine cone 661W cell line (ATCC, Virginia, USA) was cultivated in Dulbecco’s Modified Eagle’s Medium (DMEM, Gibco) complemented with 10% (v/v) fetal bovine serum (FBS, Gibco) and 1% penicillin-streptomycin (Gibco). Cells were maintained in a 37°C, 5% CO2, fully humidified incubator, and passaged twice weekly. 661W were plated at 5 x 10^4^ cells per well in a 48-well plate with a total volume 250 μL/well. 24h after plating, cells (70-90% confluency) were transfected with Lipofectamine 2000 following the manufacturer’s protocol. Briefly, sgRNA and spCas9 were mixed by pipetting 4 times, and incubated at room temperature for 5 minutes. In parallel, 2 µL of Lipofectamine™ 2000 were added to a final volume of 25μL of Opti-MEM™ Reduced Serum Medium (ThermoFisher). Then Cas9 RNP solution was added to Opti-MEM/lipid solution, vortexed for 10 seconds and incubated at room temperature for 10 minutes. 25 µL of complexed solution was added to each well to obtain a final RNP concentration of 100 nM in a total volume of 275 μL. Medium was change 24h after transfection and cells were harvested 48 hours after transfection. Cell pellets were washed with PBS and frozen at –20°C until DNA extraction.

### Subretinal injections in mice

All animal experiments were realized in accordance with the NIH Guide for Care and Use of Laboratory Animals (National Academies Press, 2011). The protocols were approved by the Local Animal Ethics Committees and conducted in accordance with Directive 2010/63/EU of the European Parliament.

C57BL/6j wild type mice (Janvier Laboratories) were used for this study. For ocular injections, mice were anesthetized by isoflurane inhalation. Pupils were dilated and sub-retinal injections of 2 µL were performed using a Nanofil syringe with a 28-gauge blunt needle (World Precision Instruments, Inc.) under an operating microscope (Leica Microsystems, Ltd.). Ophtalmic ointment (Fradexam) was applied after surgery. Eyes with extensive subretinal hemorrhage were excluded from the analysis. Optical coherence tomography (OCT) images were taken 1 month after injection under isoflurane anesthesia. Animals were euthanized by CO2 inhalation and cervical dislocation. For indels analysis, after 7 days the whole RPE and the whole neural retina were isolated without selection of transfected cells.

### Samples preparation for quantification of indels

For genomic DNA extraction, samples from each experimental condition were incubated in lysis buffer and proteinase K at 56°C overnight for retina samples, 3 hours for RPE tissue and 10min for *in vitro* cells. Further steps were performed following the manufacturer’s instructions (NucleoSpin® DNA tissue, Macherey-Nagel). We used the GXL HiFi DNA polymerase (Takara) for PCR amplification. Primers for amplifying region of interest are listed in supplementary table S3. The thermal cycler program for PCR was as follows: 98°C for 10s, followed by 60°C for 15 s and finally 68°C for 20s, with in total 30 cycles. PCR samples were then purified using following the manufacturer’s instructions (NucleoSpin® PCR and gel kit, Macherey-Nagel). Purified amplicons were verified by electrophoresis on a 1% agarose gel. For *in vivo* experiments, PCR amplicons were sent to next generation sequencing (NGS) at Massachusetts General Hospital DNA core facility. A total of 10 000 reads were generated per samples and analysis were done with a cut off of 10 reads. For *in vitro* experiments, PCR amplicons were sent to Sanger sequencing (Eurofins) and indels were analyzed using TIDE (Tracking of Indels by Decomposition) Analysis (https://tide.deskgen.com/). Analysis was done using a reference sequence from untreated samples and setting the parameters to detect a maximum indels size of 10 nucleotides.

### Statistical analysis

All statistical analyses were carried out using GraphPad PRISM version 7.0. P-values were determined by ordinary One-way ANOVA test, Dunnett’s multiple comparisons test if more than two conditions analyses or two-tailed parametric paired Student’s t-test for the analyses on base editors indels (two conditions compared). ns: non significant, * p<0.05, ** p<0.005, *** p<0.0005, **** p<0.0001.

### Immunostaining and imaging

Three days post-injection, mouse eyes were enucleated and immediately fixed in a 4% formaldehyde solution for 1 hour. Samples were either used for cryosections, flat-mount retina, or whole eye clearing and imaging.

To prepare cryosections or retinal flatmounts, eyecups were immersed in PBS- 10% sucrose for 1 hour and then PBS-30% sucrose overnight at 4°C. They were embedded in OCT medium and frozen in liquid nitrogen. 12 μm-thick vertical sections were cut with a Microm cryostat.

After 3 PBS washes, retinal flatmounts or cryosections were incubated in a blocking buffer for 1 hour and then with primary antibodies overnight at 4°C. Primary antibodies used in this study are listed in Supplementary table S4. After three washes of the sections, the secondary antibodies (Alexa Fluor 488, 594 or 647, Thermo Fischer Scientific) were added for 2 hours at room temperature, followed by three PBS washes. Retinal flatmounts or cryosections were mounted in Vectashield mounting medium (Vector Laboratories) and visualized using an Olympus confocal microscope. ImageJ software was used to process the images.

### Tissue clearing and imaging

To remove pigmentation and reduce background related to hematomas, a tissue bleaching was carried out as previously described (*31*). Then the EyeDISCO protocol was used to bleached and clear eye samples (*32*).

For whole-mount immunostaining, samples were transferred to a solution containing the primary antibodies diluted in PBSGT and placed at 37°C with agitation for 7 days. Primary antibody dilutions are described in Table S4. After six washes of 1hr in PBSGT at RT, Samples were incubated at 37°C in the secondary antibody solution for 2 days sample and washed six times during 1 hr in PBSGT at RT.

To facilitate the handling and imaging with the light-sheet microscope, tissue samples were embedded prior to clearing in 1.5% agarose (Roth,2267.4), prepared in TAE 1X (Invitrogen,15558-026).

### 3D Imaging of cleared specimens and imagine analysis

Cleared samples were imaged with a Blaze light-sheet microscope (Miltenyi Biotec) equipped with sCMOS camera 5.5MP (2560×2160 pixels) controlled by Imspector Pro 7.5.3 acquisition software.

To isolate the neural retina using the Imaris software, the surface tool was manually applied and the mask option was selected. Mask obtained with retina segmentation was used to highlight staining of the neural retina (using Recoverin).

### SD-OCT Imaging

Spectral-domain optical coherence tomography (SD-OCT) was performed 1 month post injections. The mice’s pupils were dilated with tropicamide (Mydriaticum, Théa) and phenylephrine (Néosynephrine, Europhta). The animals were anesthetized by inhalation of isoflurane (Isorane, Axience) and placed in front of the SD-OCT imaging device (Bioptgen 840 nm HHP; Bioptgen). The eyes were kept moist with 9% NaCl during the whole procedure. Image acquisitions were performed using the following parameters: rectangular scan/1000 A-scan per B-scan/100 B-scan 1 frame. ImageJ software was used to process the images as .avi.

## Supporting information

Supplementary results

## Acknowledgements

We are grateful to S. Giovagnoli, A. Saint-Julien, Y. Rasool, S. Labou and H. Eriksson for preliminary in vitro work, samples preparation and indel analysis, Q. Cesar and R. Goulet (phenotypic facility) for OCT and micron recordings, Jean Baudry for DLS access at Chimie Paris Tech.

## Author Contributions

CB, JP and DD designed experiments. ADC and MA produced Cas9 proteins. CB, JP, PO, CI and BS optimized TEM analysis. JP and PO performed and analyzed DLS recordings. CB designed sgRNAs and performed transfection on cell lines. CB and JP performed sub-retinal injections in mice. CB, JP, CR, HM and PO dissected mice eyes, prepared and analyzed NGS samples. JP and CB performed histology, immunostaining and imaging. SF performed clearing and 3D imaging. JP optimized and analyzed OCT recordings. HM performed the experiment on ABE. DR performed and DR and SF analyzed molecular biology experiments in relation with inflammation. CB, JP and DD designed the study and wrote the manuscript. AE, JPC and SF provided scientific input and gave feedback on the manuscript.

## Data and materials availability

The data that support the findings of current study are available from the corresponding author upon reasonable request.

