## Supplementary results for "Direct delivery of Cas9 or base editor protein and guide RNA complex enables genome editing in the retina"

**(See additional movies)**

**Figure S1: Video of a 3D rendering of a mouse eye injected with Cas9 RNPs.**

Eyes were collected three days after 2 µL subretinal injection of buffer (control) or 30 µM Cas9 RNPs. Whole-mount immunolabeling for the photoreceptors (Recoverin, red) and Cas9 protein (green). Neural retina was isolate after manual segmentation.

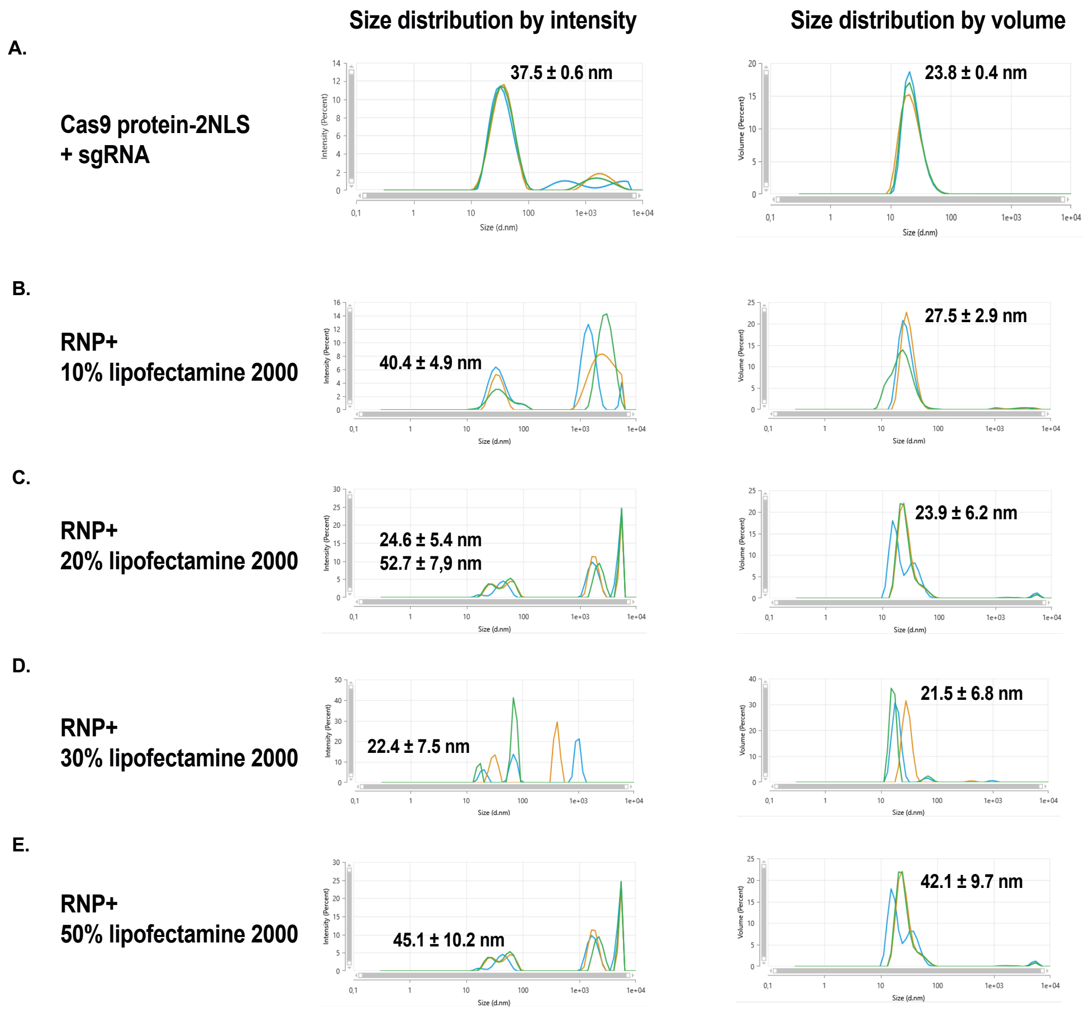


**Figure S2: Size determination of Cas9 RNP and Cas9 RNP complexed with lipofectamine 2000.**

Size distribution by intensity and volume of Cas9 protein and its sgRNA: (A) without complexation or (B to E) complexed with an increase volume of lipofectamine 2000 (lipids represent from 10% to 50% of total volume of injection). Three different measures (in blue, red and green) were performed at 5-minute intervals.

A.

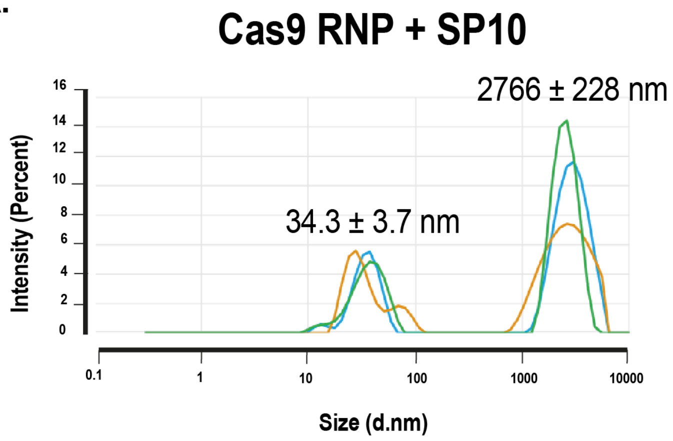


B.

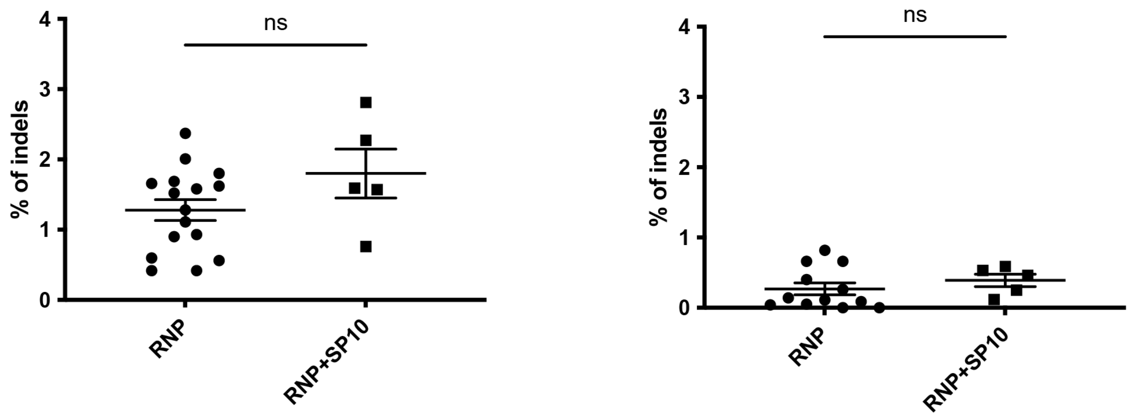


**Figure S3: Mixing Cas9 RNP with SP10 doesn't improve the editing efficiency**

**A. Size determination of Cas9 RNP with SP10 peptide.** Size distribution by intensity and volume of Cas9 protein and its sgRNA: (A) without complexation or (B to E) complexed with an increase volume of lipofectamine 2000 (lipids represent from 10% to 50% of total volume of injection). Three different measures (in blue, red and green) were performed at 5-minute intervals

**B.** Indels in the RPE and neural retina after sub-retinal injection of Cas9 RNPs mixed or not with SP10. NGS analysis was performed 7 days after injection. Mean ± SEM. Student's t-test. ns= non-significant.

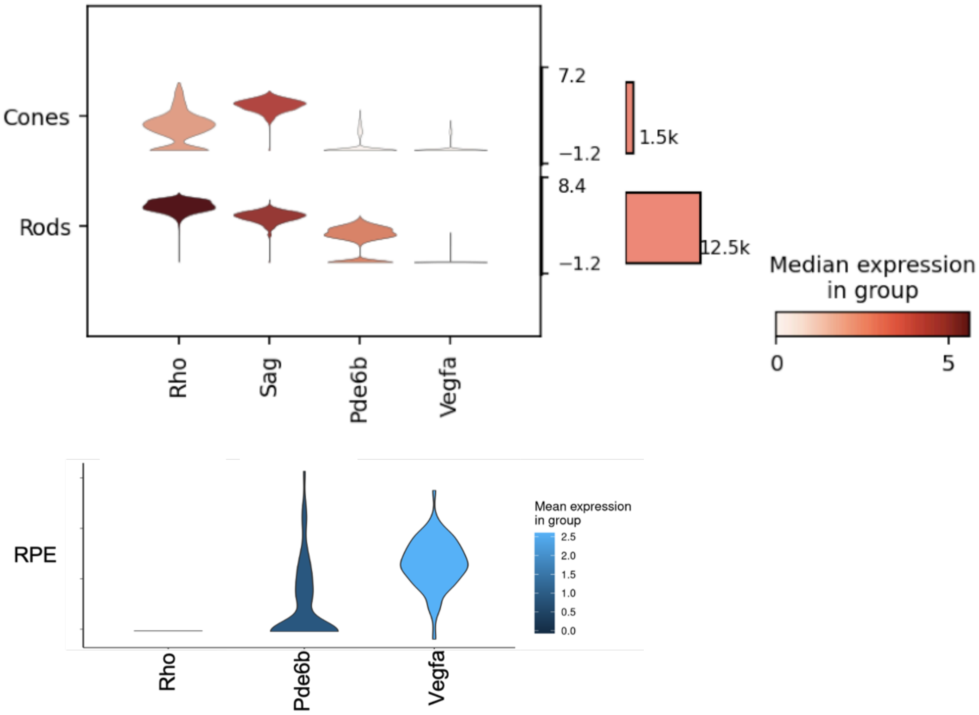


**Figure S4: Expression level of *Vegfa*, *Pde6b*, *Sag* and *Rho* gene in cones, rods and RPE of wild-type adult mice.**

Upper plots, RNAseq analysis based on wild type adult mice database (GSE132229) (*31*)

Lower pots, RNAseq analysis based on wild type adult mice database from (*32*). Sag was not detected in this experiment.

| Cas9 protein | Concentration and volume | Molecules |
|---|---|---|
| ***In vitro*** **(5.10^5 cells)** | **100 nM in 275 µL** | **1.7 10^13** |
| ***In vivo*** **(7.10^9 neural retina cells)** | **30 µM in 2 µL** | **3.6 10^13** |

**Table S1: Calculation of Cas9 molecules transfected *in vitro* or injected in the sub-retinal space *in vivo*.**

*In vitro*, 5.10^5 were transfected by 100nM in a total volume of 275 µL. *In vivo*, 2 µL of 30 µM Cas9 RNPs solution were sub-retinally injected. Neural retina cells number is estimated at 7.10^9 cells (Jeon et al. 1998). Number of Cas9 molecules is determined based on Avogadro constant (1 Na= 6.02 10^23 $mol^{-1}$).

| Gene (exon) targeted | sgRNA name | Target sequence |
|---|---|---|
| ***Vegfa*** **(exon3)** | Vegfa sgRNA | CTCCTGGAAGATGTCCACCA |
| | Rho sgRNA 1 | GGTTCCGCCAGGTAGTACTG |
| | Rho sgRNA 2 | TAGAGCGTGAGGAAGTTGAT |
| ***Rho*** **(exon 1)** | Rho sgRNA 3 | GCCAGCATGGAGAACTGCCA |
| | Rho sgRNA 4 | CAAGACGGAGACCAGCCAGG |
| ***Rho*** **(exon 5)** | Rho sgRNA 5 | GAGGCGTCGTCATCTCCCAG |
| | Pde6b sgRNA 1 | GCCGTGGCGCCAGTTGTGGT |
| | Pde6b sgRNA 2 | GAGTAGGGTAAACATGGTCT |
| ***Pde6b*** **(exon 13)** | Pde6b sgRNA 3 | GTGGTAGGTGATTCTTCGAT |
| | Sag sgRNA 1 | TCCCTTGGGATGTACGTCAG |
| ***Sag*** **(exon 2)** | Sag sgRNA 2 | GACCTTCTTGAAGATGACGT |
| ***Sag*** **(exon 8)** | Sag sgRNA 3 | AGACATGAAGAACTGCCAGG |

**Table S2: sgRNA sequences used in this study.**

| Sanger Primers | | Sequence |
|---|---|---|
| *Vegfa* exon 3 | Fw | CAAATCTGGGTGGCGATAGA |
| | Rev | AGATGGTCAAATCGTGGAGAG |
| *Rho* exon 1 | Fw | ATATCTCGCGGATGCTGAAT |
| | Rev | GCAAAGAAGCCCTCGAGATT |
| *Rho* exon 5 | Fw | CATGTGGACTTCGGTTTTCC |
| | Rev | GGAGCCTCATTTTGCTTTCA |
| *Pde* exon 13 | Fw | GGCCAGTGAGAACAAGGAAC |
| | Rev | GCTCCAGAAGGCAGTGGTTA |
| *Sag* exon 2 | Fw | GCTGCACACTACAGTAAAGTACAACTG |
| | Rev | TCCACTGTTGTTCCTTGGTTC |
| *Sag* exon 8 | Fw | GGTATCTGAAACCATGACACAGA |
| | Rev | GGGGAGAGCCAACAAGTACA |
| **NGS Primers** | | **Sequence** |
| *Vegfa* exon 3 | Fw | TCATGGATGTCTACCAGCGA |
| | Rev | AATGCCCCTCCTTGTACCAC |
| *Sag* exon 1 | Fw | GAGGAAGTCACAGGAGTGGG |
| | Rev | GCCTCTCTTCTGTGCCACTTA |
| *Sag* exon 8 | Fw | GCTCTGTGCGGTTACTGATC |
| | Rev | AGTGGAAATAAGCTGGTAATTTGCA |

**Table S3: Primers used to amplify the targeted regions of each gene for Sanger or NGS sequencing.**

| Antigen | Species | Dilution | Source |
|---|---|---|---|
| Iba1 | Rabbit polyclonal | 1/250 | WAKO |
| Recoverin | Rabbit polyclonal | 1/3000 | Millipore MAB 5585 |
| SpCAS9 | Mouse monoclonal | 1/1000 | Genscript A01935-40 |

**Table S4: Primary antibodies used for immunostaining.**
